# Perceived risk of forward versus backward balance disturbances while walking in young and older adults

**DOI:** 10.1101/2025.01.08.632059

**Authors:** Shreya Jain, Nicolas Schweighofer, James M. Finley

## Abstract

Falls, which often result from trips or slips, pose a major health concern, particularly among older adults. Experiencing falls or near-falls from balance disturbances can shape subsequent gait-related decisions, as individuals may avoid situations they perceive as risky. Here, we explore whether perceptions of risk are sensitive to the direction of previously experienced balance disturbances – forward or backward – and whether these perceptions change with age. Twenty young and twenty older adults walked on a split-belt treadmill while performing a two-alternative forced-choice task, where they indicated their preference between a forward-falling trip and a backward-falling slip. We varied the perturbation magnitudes using an adaptive staircase algorithm to obtain multiple trip-slip equivalence points. Using a mixed-effects linear model, we estimated the slope of the trip-slip relationship, which quantified the direction and strength of the sensitivity of perceived risk to perturbation type. To assess reliability, we repeated the procedure on a second day. Additionally, we investigated two potential reasons underlying any observed sensitivity – 1) emotional responses measured by state anxiety, and 2) physical responses measured by peak center of mass velocity. We found that both young and older adults perceived slips to be riskier than trips, with no group difference in sensitivity. The relative sensitivity to slips versus trips was moderately reliable across two days of testing, though most participants were less sensitive to perturbation direction on the second day. Neither state anxiety responses nor peak center of mass (CoM) velocity explained the directional sensitivity, though CoM velocity was higher during slips than trips for both age groups. These results suggest that the characteristics of experienced balance disturbances may influence behavioral decisions. Objectively measuring individual differences in the perceived risk of balance disturbances and the ability to recover from these disturbances, can potentially improve fall risk assessment and inform personalized training interventions.

## Introduction

Falls are a public health concern, particularly among older adults for whom they can lead to devastating injuries and significant reductions in quality of life [1,2]. Falls result from several risk factors, which include physical ability, features of the environment, and individual differences in risk-taking behaviors [3–6]. The decision to engage in potentially risky behaviors, such as walking in crowded or cluttered areas, relies on one’s perceived risk of losing balance, falling, or injuring oneself in the current environment, and this perception may be informed by previous experiences in the same or similar environments. Falls most commonly occur during walking, precipitated by losses of balance such as trips and slips [7–9]. While trips and slips are commonly categorized together as a cause of falls, experiencing these two losses of balance may influence future gait behaviors differently [10]. Therefore, it is important to understand how these balance disturbances are perceived relative to each other and how their associated perceived risk informs decisions about future behaviors.

Risk assessment largely relies on the possible outcomes that may occur in a given situation and their perceived severity [11,12]. In the context of walking in risky environments, trips and slips are examples of possible outcomes. Trips commonly occur when the foot collides with an obstacle, obstructing forward motion and causing a loss of balance in the forward direction [13]. Slips occur when there is insufficient friction between the foot and the walking surface, leading to a loss of balance in the backward direction [14]. When faced with walking options that may result in slips or trips, people’s choices may be influenced by how they perceive the relative riskiness of these perturbations. Risk perception may be influenced by perceived changes in body kinematics as well as emotional responses to experienced balance disturbances. Slip-like perturbations generated on a split-belt treadmill cause quicker stepping responses and larger whole-body center of mass (CoM) deviations than trip-like perturbations, suggesting that it may be more difficult to recover from slips [15]. This greater challenge may cause slips to be perceived as being more severe and riskier than trips. Emotions such as fear and anxiety are important determinants of perceived risk and have also been shown to influence perceptions of self-motion under postural threat (e.g. standing on an elevated platform) [12,16–18]. Therefore, if trips and slips are perceived differently in terms of their severity, this may be due to the emotional responses that they evoke.

Older adults are often more risk-averse than young adults in several domains of decision-making [19–21]. In situations concerning health, safety, and recreational activities, older adults report perceiving more risk than young adults [22]. While there is some evidence to suggest that these same trends are observed in motor decisions related to reaching movements [23], their extension to gait and balance related decisions remains to be seen. If older adults are sensitive to the direction of balance disturbances such that they perceive them to be differently risky, their increased risk aversion may lead to this sensitivity being heightened compared to young adults.

The objective of this study was to assess the relative perceived risk of forward versus backward balance disturbances during walking, herein described as trips and slips, respectively, in older and younger adults by determining equivalence points between the two types of perturbations [24,25]. For example, if a large trip is perceived to be equivalent to a small slip, this suggests that the slip is perceived to be a worse outcome than a trip. We hypothesized that backward-falling slips would be perceived to be riskier than forward-falling trips, and this sensitivity to perturbation direction would be greater in older adults than young adults. A secondary objective of this study was to investigate if changes in whole-body CoM kinematics or measures of state anxiety explained the observed differences in perceived risk between the two directions of perturbations. Based on previous work showing larger CoM responses to slips than trips, we hypothesized that differences in CoM responses between trips and slips would correlate with any observed sensitivity in perceived risk. Lastly, since anxiety influence risk perception and postural control in threatening conditions, we hypothesized that differences in self-reported anxiety in response to trips and slips would correlate with relative perceived risk between the two perturbation types.

## Methods

### Experimental Procedures

Twenty young adults (11 female, 27 ± 5 years) and twenty older adults (12 female, 71 ± 4 years) participated in this study over two separate visits, at least a week apart. All participants provided their informed consent in accordance with a protocol approved by the Institutional Review Board of the University of Southern California. All participants indicated they could walk without an assistive device for 30 minutes and did not have any recent injuries that prevented them from doing so.

We began the first visit by having participants complete a set of clinical assessments.

We assessed overall balance ability using the MiniBESTest [26]. The assessment consists of 14 items with a maximum possible score of 28, divided into four subscales that assess anticipatory postural control, reactive postural control, sensory orientation, and stability during gait. This was followed by the Activities-specific Balance Confidence (ABC) scale [27] as a measure of perceived balance ability, and a History of Falls questionnaire [9] to record falls that occurred in the 12 months preceding the visit. To estimate physical activity levels, participants provided self-reports of the approximate time in minutes per week that they engage in exercise.

### Self-Selected Walking Speed

We determined participants’ self-selected walking speed for both overground and treadmill walking. Overground walking speed was determined using an average of three 10- meter walks. To determine self-selected walking speed on the treadmill (Fully Instrumented Treadmill, Bertec Corporation, Columbus, OH, USA), we used an adaptive staircase algorithm [28]. Starting at 80% of the overground speed, treadmill speed was first increased by steps of 0.1 m/s for 10 s until the participant verbally indicated that they were at their comfortable walking speed. Next, the speed was further increased by 0.1 m/s, followed by a down-ramp with a step size of 0.1 m/s until the participant verbally indicated that they were at their preferred walking speed. The up-ramp and down-ramp were repeated a second time with a step size of 0.05 m/s. The final self-selected walking speed was computed as the average of the four indicated preferred walking speeds. This procedure was followed by an acclimation trial with two minutes of walking at the determined self-selected walking speed.

### Familiarization with Perturbations

To determine differences in perceived risk of forward versus backward balance disturbances, we accelerated or decelerated the belts of a split-belt treadmill while participants walked [15,28,29]. Perturbations were delivered by rapidly changing the speed of one belt during the swing phase so that the belt speed was faster or slower than the self-selected speed at the subsequent foot-strike [28]. Accelerations led to forward-falling perturbations, and decelerations led to backward-falling perturbations. We will refer to these as trips and slips, respectively, while acknowledging that changes in treadmill speed may not precisely recreate the mechanical stimuli of real-world trips and slips [30–32]. While handlebars were attached at the front of the treadmill, participants were verbally discouraged from using them for external support.

Participants underwent a short familiarization procedure to attenuate any effects of the novelty of split-belt treadmill perturbations on choices. This included two familiarization blocks, one with trips and one with slips. One familiarization block consisted of two minutes of walking with two perturbations, each with a magnitude of 0.6 m/s. After each block, participants responded to a State Anxiety Scale, which was used to assess their emotional response to both directions of perturbations [33]. This allowed us to test if differences in the anxiety response could explain any observed differences in the perceived risk of trips and slips. The State Anxiety Scale consisted of 9 items related to somatic anxiety (5 items), concentration disruption (2 items), and worry (2 items), which were scored on a 9-point Likert scale, ranging from 1 (*‘I do not feel this at all’*) to 9 (*‘I feel this extremely’*).

### Two-Alternative Forced Choice Task

To quantify differences in perceived risk based on perturbation direction, we used a two-alternative forced-choice task wherein participants experienced one trip and one slip and then responded to the prompt - “If you had to repeat one of these perturbations, which one would you choose?” The prompt appeared on a monitor placed at the front of the treadmill, and participants indicated their choice using a keypad (Fig 1). Because there was no incentive provided for choosing the worse outcome, we assumed that this question prompted participants to choose the perturbation that was perceived to be less risky. We verified this assumption by asking participants to verbalize their decision-making process at the end of the study.

**Fig 1.**
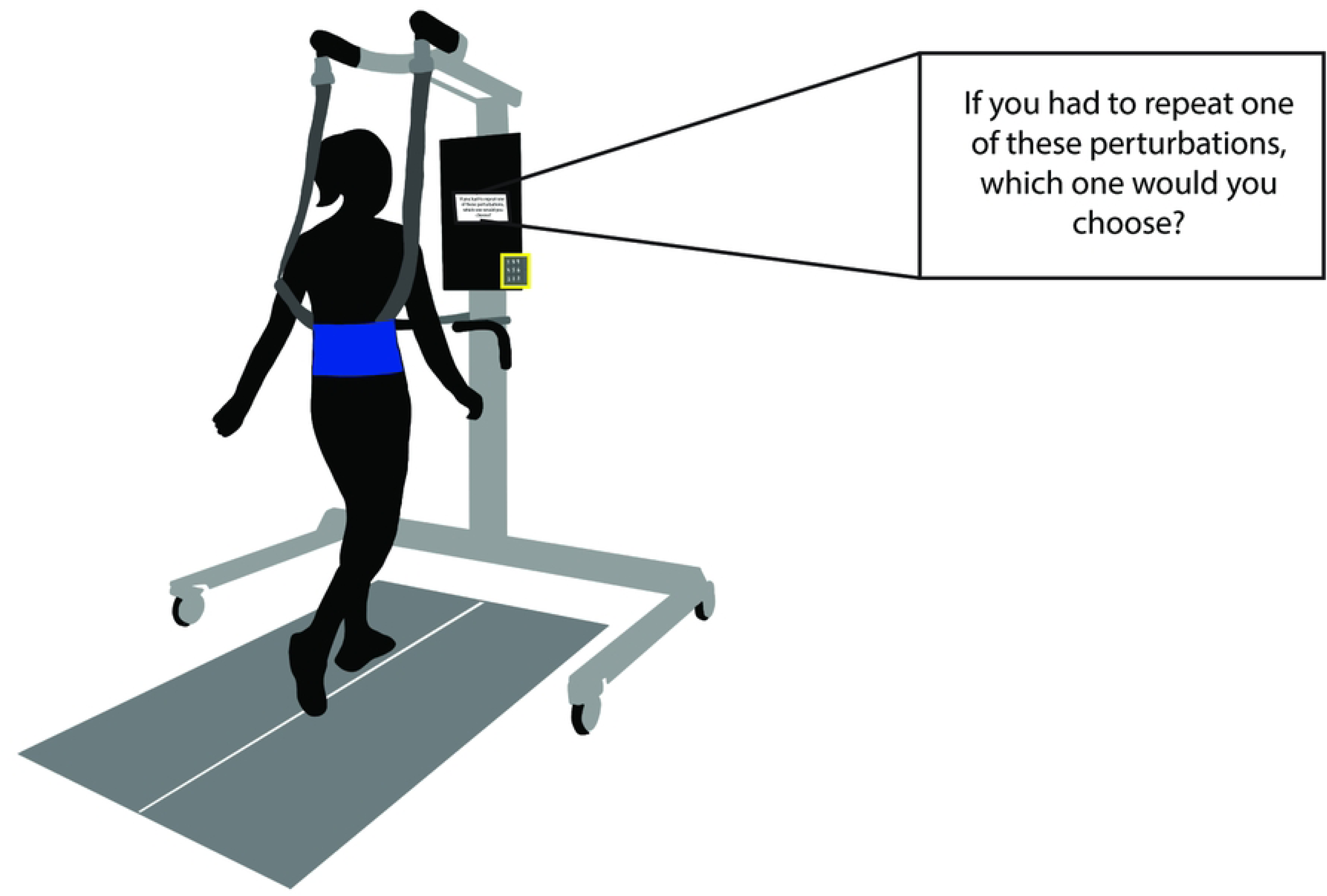
Experimental setup. Participants walked on a split-belt treadmill and received tripping and slipping perturbations. The decision prompt (enlarged on the top right) appeared on a monitor in front of the treadmill, and participants indicated their choices using a keypad (highlighted in the yellow box).

We used seven different trip magnitudes ranging from 0.3 to 0.6 m/s in increments of 0.05 m/s. For each trip magnitude, an adaptive staircase procedure was used to determine the equivalent preferred slip magnitude, referred to here as the equivalence point [24,25]. The staircase started with a trip and a slip of equal size and consisted of two up-ramps and two down-ramps. In the up-ramps, the size of the slips was increased until a trip was chosen, and this was defined as an inflection point. Next, the slip size was increased once more before beginning the down-ramps. In the down-ramps, the size of the slips was decreased until a slip was chosen. The step size for slip magnitude changes was 0.1 m/s in the first two ramps and 0.05 m/s in the second two ramps. An example of the staircase procedure for a trip magnitude of 0.4 m/s is shown in Fig 2. The equivalence point was computed as the average of the four inflection points.

**Fig 2.**
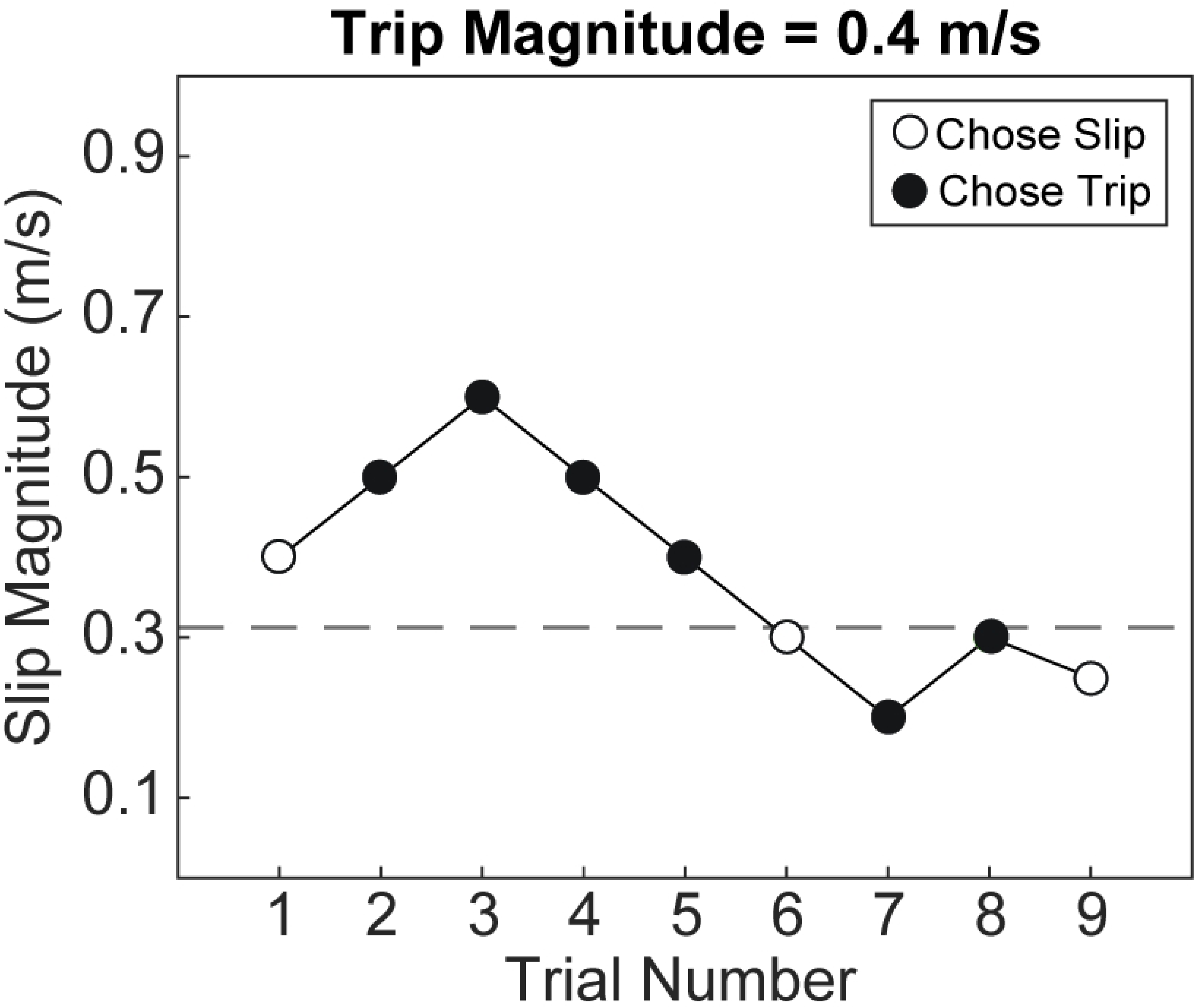
Example of the staircase procedure for a trip magnitude of 0.4m/s. At trial 1, both the trip and slip were of the same magnitude. Open circles indicate a choice of the slip perturbation, and filled circles indicate a choice of the trip perturbation. For each up-ramp, the inflection point is the trial at which they chose the trip. For each down-ramp, it is when they chose the slip. Inflection points occurred during trials 2, 6, 7, and 9, and the equivalence point was computed as the average of the four inflection points. In this example, the equivalence point is 0.31, implying that a slip of 0.31m/s is perceived to be equivalent to a trip of 0.4m/s.

This procedure was repeated on both the left and right sides, yielding 14 equivalence points between trips and slips for each participant. To assess the reliability of this procedure and the stability of the equivalence points, we repeated this procedure on a second visit at least a week after the first one. At the end of the second visit, participants provided a subjective description of their decision-making strategy.

### Data Acquisition

While participants walked, positions of the pelvis and feet were recorded with an 11- camera Qualisys Oqus camera system (Qualisys AB, Göteborg, Sweden), using a configuration of thirteen reflective markers – one on each greater trochanter, three on the pelvis (posterior superior iliac spines, lumbosacral joint) and four on each foot (fifth metatarsal, lateral malleolus, first toe, heel). These data were available for 18 young adults and 19 older adults.

### Data Processing

We post-processed the kinematic data in Visual3D (C-Motion, Rockville, MD, USA) and MATLAB R2022b (MathWorks, USA). Marker positions were low-pass filtered using a 4th order Butterworth filter with a cutoff frequency of 6 Hz. Heel strike and toe off events were identified as the peak anterior position of the heel marker and peak posterior position of the toe marker, respectively. We estimated the center of mass of the pelvis in Visual3D with the default model of the pelvis and we used this as a surrogate for the whole-body center of mass (CoM) [34]. CoM velocity was computed as the first derivative of CoM position.

Previous work suggests that CoM velocity may be an important sensory signal for perception of treadmill perturbations [35]. Therefore, to test whether the change in CoM velocity during the perturbations may explain inter-individual differences in the preference between the two perturbation directions, we analyzed the peak CoM velocity during the first trip-slip pair of each staircase. We chose the first pair because both perturbations were of equal size at the beginning of our staircase algorithm. This yielded 14 total perturbations in each direction per day of testing. We first computed the resultant CoM velocity during each step as the square root of the sum of the squares of the velocities in the anterior-posterior, mediolateral, and vertical directions. Anterior-posterior velocity was computed relative to the treadmill belt. We then identified the peak forward and backward CoM velocity during the trips and slips, respectively (Fig 3A). We also computed peak forward and backward CoM velocities for 14 steps of unperturbed walking from the baseline trial. Next, we characterized the effect of the trips on the CoM as the magnitude of the difference between the average peak CoM velocity during all 14 trips and 14 unperturbed steps (Fig 3B). Similarly, for slips, we computed the magnitude of the difference between the average peak negative CoM velocity during slips and unperturbed steps.

**Fig 3.**
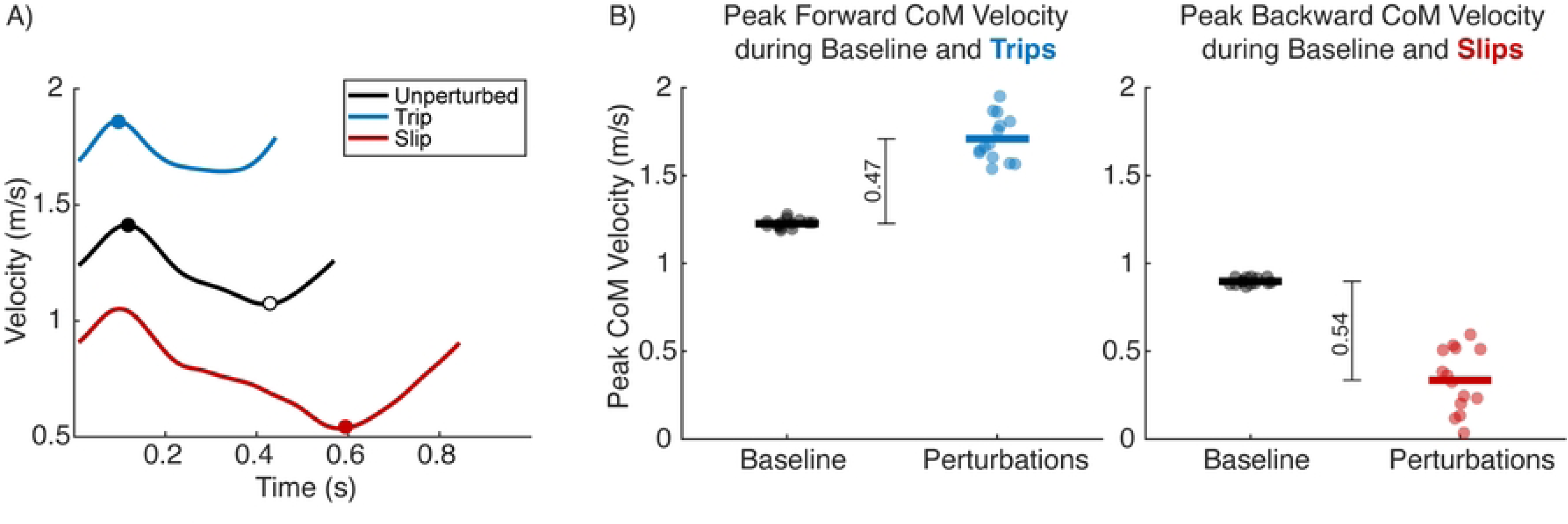
Example of the calculation of CoM velocity responses to perturbations. A) Example of anterior-posterior CoM velocity during an unperturbed step (black), a trip (blue), and a slip (red). The circles indicate the peak forward and backward CoM velocities. B) Example of peak forward resultant CoM velocities (left panel) during unperturbed and trip steps, and peak backward CoM velocities (right panel) during unperturbed and slip steps. Each point represents one step. Horizontal lines represent mean values. The overall CoM response to trips was computed as the difference between the mean peak forward CoM velocity during trips and unperturbed walking (black vertical line, left panel). The overall response to slips was computed as the magnitude of the difference between the mean peak backward CoM velocity during slips and unperturbed walking (black vertical line, right panel).

### Statistical Analysis

We used t-tests to test for potential differences in walking speed, MiniBESTest, ABC scores, and minutes of exercise per week between the young and older adults. In case of non-normality in the data, we used the Mann-Whitney U-test for independent samples. Normality was assessed using the Shapiro-Wilk test.

To quantify sensitivity to perturbation direction, we fit a linear mixed effects model with preferred slip magnitude as the dependent variable, trip magnitude as the independent variable, an interaction between trip magnitude and age group, and random slopes for participants (Equation 1).

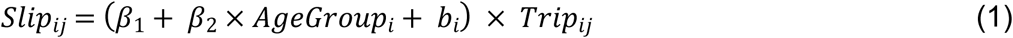

Here, *i* refers to the participant number, *j* refers to the staircase number, AgeGroup is 0 for young and 1 for older adults. β_1_ and β_2_ represent the regression coefficients for the trip magnitude and the interaction term between trip and age group, respectively. *b_i_* represents the random effect of participants.

The intercept of this model was set to 0, as a non-zero intercept here implies a preference for some slip magnitude even in the absence of a trip, which is not a meaningful or plausible interpretation. Because including a main effect of age group would yield a non-zero intercept, only the interaction term between trip magnitude and age group was included in the model. The term in the parentheses in Equation 1 then represents the slope of the relationship between trips and subjectively equivalent slips, which indicates the direction and strength of the preference for a perturbation direction. The final slope for each participant was calculated as a sum of the fixed and random effects. Slopes less than 1 imply a preference for trips, slopes greater than 1 imply a preference for slips, and a slope equal to 1 implies indifference between the two directions.

To determine the reliability of these slopes as measures of sensitivity to perturbation direction, we fit this model separately to data from a second day of testing. We computed the intra-class correlation coefficient (ICC) between the individual slopes obtained on the two days as a measure of the agreement between them. Additionally, to test for the presence of a bias in slopes on Day 2 relative to Day 1, we included testing day as a covariate in the mixed effects model described above (Equation 2).

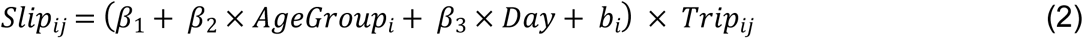

The ages of our older adult participants had a wide range, with the lowest being 66 and the oldest being 84 years of age. To account for potential age-dependent differences within this group that may not be captured when grouping them together, we performed an exploratory analysis with the older adults wherein we replaced the age group variable in equation 1 with age as a continuous variable. This allowed us to test if the older adults on the higher end of the age range in this group differed from those on the lower end. We performed this analysis separately with data from the two days of testing.

To investigate the influence of State Anxiety responses to perturbations on people’s preferences for slips versus trips, we first determined if these responses differed between the two perturbation types and age groups. Because anxiety scores were aggregated Likert scale responses, we used a generalized linear model with a quasi-Poisson error distribution to account for the skew and overdispersion observed in this data. This model included the state anxiety scores as the response variable and perturbation type, age group, and the interaction between them as the predictors. Next, we applied a linear mixed effects model similar to the ones described above in equations 1 and 2 and included the difference in total state anxiety scores between slips and trips as a covariate (Equation 3). Because state anxiety was only assessed on Day 1, this model only included data from this day.

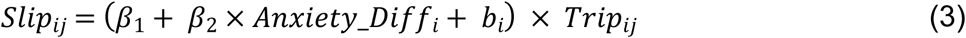

Lastly, we analyzed the CoM velocity responses to trips and slips to determine if the difference in these responses between slips and trips may influence the observed preferences. First, we used a linear model to test for an effect of perturbation type, age group, testing day, and their interactions on changes in peak CoM velocity relative to baseline. Next, we applied a linear mixed effects model similar to that described above in equation 3, and included the difference between CoM velocity responses to slips and trips as a covariate (Equation 4). We applied this model separately to data from the two days of testing.

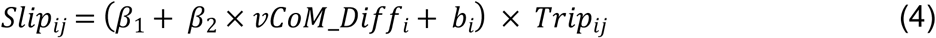

All statistical analyses were performed using R (version 4.3.1) within RStudio (version 2023.03.0+386).

## Results

Our clinical assessments revealed some aspects of physical capacity and activity that were dissimilar between our young and older adults, while others were similar (Table 1). For example, our older adults had lower MiniBESTest scores than the young adults (Mann-Whitney U-statistic = 370, p < 0.001). The number of participants who experienced a fall in the 12 months preceding the study was also higher among older adults (n=10) than young adults (n=7). However, while all falls among the young adults occurred during sports-related activities, 6 out of 10 older adult fallers reported experiencing falls during activities of daily living (e.g., walking on a sidewalk, navigating stairs, walking a dog). Despite the presence of between-group differences in balance and falls, there were no between-group differences in walking speed (p=0.122), balance confidence scores (p=0.284), or physical activity (p=0.100).

**Table 1:** Participant characteristics.

|  | Young Adults (n = 20) | Older Adults (n = 20) | p-value |
| --- | --- | --- | --- |
| Age (Years; Mean ± SD) | 27 ± 5 | 71 ± 4 | - |
| Sex | 11F/9M | 12F/8M | - |
| Treadmill Walking Speed (m/s; Mean ± SD) | 1.02 ± 0.13 | 1.09 ± 0.15 | 0.122 |
| Number of Fallers | 7 | 10 | - |
| MiniBESTest Total Score (Median (IQR)) | 28 (IQR 27-28) | 26 (IQR 25-26) | < 0.001 |
| ABC Scale Score (Median (IQR)) | 96 (IQR 90-100) | 95 (IQR 87-97) | 0.284 |
| Weekly physical activity (minutes; Mean ± SD) | 284 ± 216 | 427 ± 296 | 0.100 |

We quantified differences in the sensitivity to perturbation direction by estimating the slopes of a linear model relating trip magnitude to the subjectively equivalent slip. Slopes less than 1 implied a preference for trips, slopes greater than 1 implied a preference for slips, and slopes equal to 1 implied indifference to perturbation direction (Fig 4A). We found a significant association between trip magnitude and the equally preferred slip on both days, with slopes generally less than 1, indicating a preference for trips over slips (Day 1: β = 0.88, SE = 0.04, p < 0.001; Day 2: β = 0.96, SE = 0.04, p < 0.001). There was no significant interaction between trip size and age group, suggesting that the strength of the sensitivity to perturbation direction was not different between young and older adults (Day 1: p=0.347; Day 2: p=0.133). In assessing the variance explained by our model, the marginal R^2^ (representing variance explained by the fixed effects only) on days 1 and 2 was 0.32 and 0.34, respectively. The conditional R^2^ (representing variance explained by both fixed and random effects) on the two days was 0.58 and 0.61, respectively. These results suggest that overall, our sample of young and older adults was sensitive to perturbation direction such that they perceived trips to be less severe than slips, with no difference in sensitivity between age groups. Participants’ subjective responses indicated that they chose perturbations that felt “easier to recover from” (n=16) or felt less intense” (n=7). Several young and older adults reported perceiving the slips to be worse and choosing them only when the trip magnitude was much greater than the slip (n=16).

**Fig 4.**
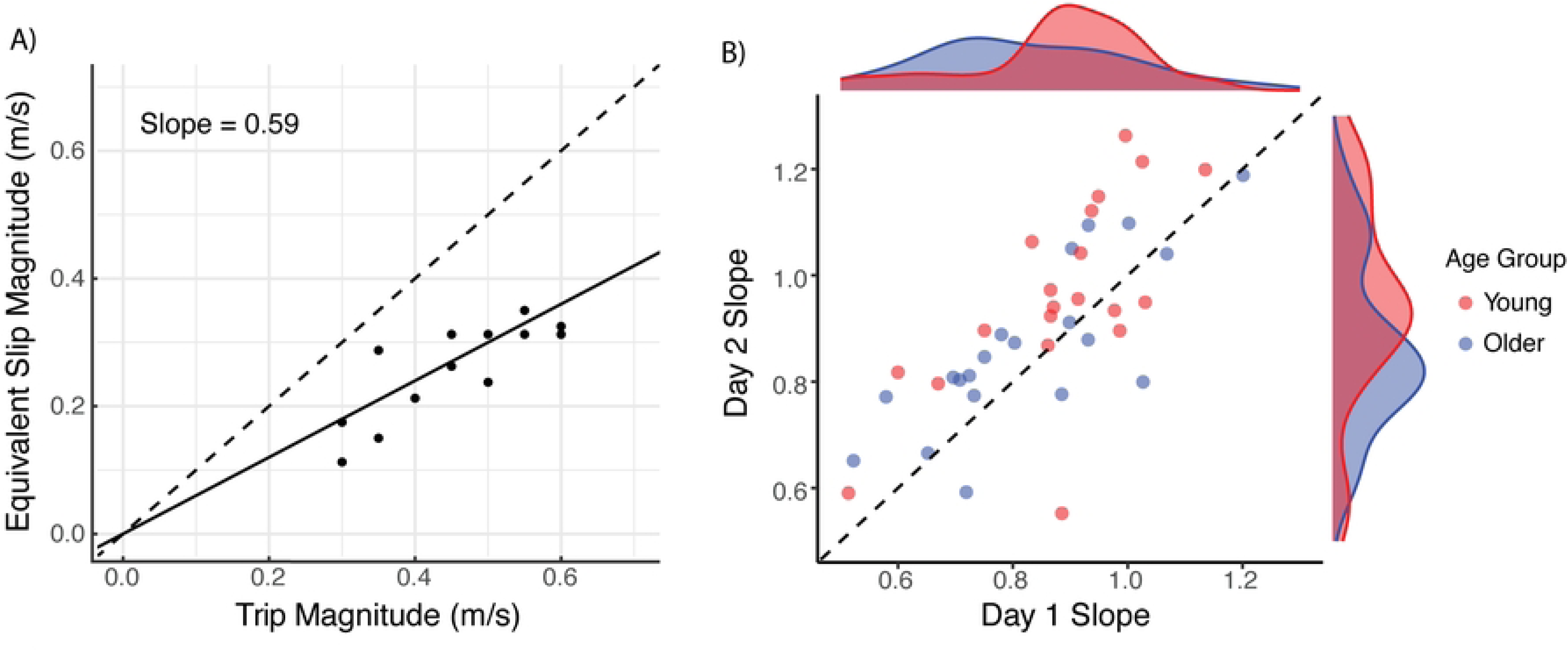
Scatterplots of slip-trip equivalence points. A) Scatterplot of equivalence points for one representative participant during a single visit. The black dashed line is the unity line, and the black solid line is the linear fit from the mixed effects model. For this participant, all equivalence points lie below the unity line, and the slope is less than one, indicating that trips were preferred to slips. B) Scatterplot of slopes obtained on Day 1 and Day 2 for all participants. Each point represents one participant, with red indicating young and blue indicating older adults. The black dashed line is the unity line. The density plots show the distribution of the slopes on Day 1 (x-axis) and Day 2 (y-axis) for both young (red) and older (blue) adults.

We next examined the stability of people’s preferences for trips versus trips by repeating the staircase procedure on a second day and comparing the model slopes on day 2 to those observed on day 1 (Fig 4B). On day 1, 17 out of 20 young adults and 16 out of 20 older adults had slopes less than 1 (young adults = 0.88 ± 0.15; older adults = 0.83 ± 0.17). On day 2, 13 out of 20 young adults and 15 out of 20 older adults had slopes less than 1 (young adults = 0.96 ± 0.19; older adults = 0.87 ± 0.16). The two-way ICC coefficient was 0.69 (95% confidence interval = 0.45-0.83), implying moderate reliability of model slopes across two days of testing. There was a significant interaction effect of trip magnitude and testing day on preferred slip magnitudes, such that model slopes were greater on day 2 than day 1 (β = 0.06, p < 0.001). These results suggest a reduction in sensitivity to perturbation direction on day 2, with slopes increasing towards 1.

To account for the wide age range among the older adults, we performed an exploratory analysis by treating age as a continuous variable in equation 1. However, on both days of testing, there was no significant interaction effect between trip magnitude and age on the preferred slip magnitude (Day 1: p = 0.31; Day 2: p = 0.97). Therefore, there did not appear to be any differences in the preference between forward and backward balance disturbances between the older adults on the lower and the higher end of the age range in our sample.

Next, we evaluated if differences in the emotional response to trips and slips as measured by the State Anxiety Scale might explain people’s preferences. Because this was a 9- item, 9-point Likert scale, the scores were between 9 and 81. There was no effect of perturbation direction (p=0.774), age group (p=0.198), or their interaction (p=0.417) on state anxiety scores (Fig 5A). There was also no interaction effect of trip magnitude and the difference in state anxiety response to slips and trips on the preferred slip magnitudes on day 1 (p=0.345). Thus, people’s preferences were not explained by differences in self-reported anxiety.

**Fig 5.**
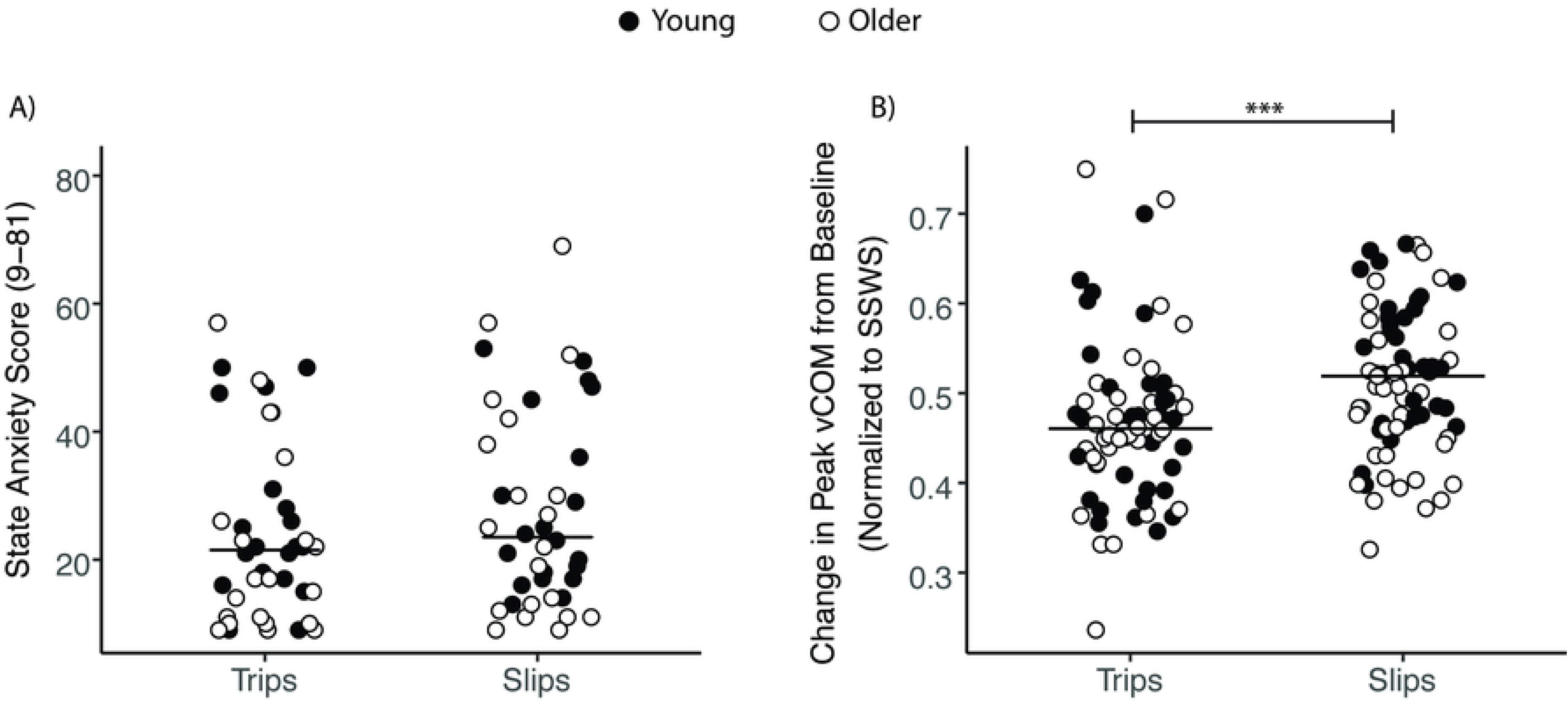
State anxiety and CoM velocity responses to trips and slips. A) Scatterplot of State Anxiety scores in response to trips and slips for young (filled circles) and older (open circles) adults. Horizontal lines represent group medians. B) Scatterplot of change in peak CoM velocity from baseline in response to trips and slips, normalized to self-selected walking speed (SSWS), for young (filled circles) and older (open circles) adults. Change in peak CoM velocity from baseline walking was higher in response to slips than trips (***p < 0.001), with no difference between age groups.

Lastly, we evaluated if differences in the biomechanical effects of slips versus trips could explain inter-individual differences in their preferences. Overall, slips led to greater whole-body disturbances than trips, as measured by the changes in CoM velocity relative to unperturbed walking (Fig 5B). We found a main effect of perturbation direction on peak CoM velocity relative to baseline, indicating that there was a larger change in peak CoM velocity in response to slips than trips (β = 0.08, p < 0.001). There was no between-group difference in changes in peak CoM velocity (p=0.914), no main effect of testing day (p=0.715), and no interaction between these variables. There was also no interaction effect of trip magnitude and the difference between peak CoM velocity between trips and slips on the preferred slip magnitudes on both day 1 (p=0.860) and day 2 (p=0.636).

## Discussion

When we experience losses of balance while walking, these experiences impact subsequent decisions about how and where we walk. However, little is known about how we process the precise characteristics of experienced losses of balance and if this evaluation process varies in older age when fall-related injuries may have more devastating consequences. As trips and slips during walking are the most common causes of falls in both young and older adults, we used a two-alternative, forced-choice procedure to determine how people evaluated the relative risk between forward-falling trips and backward-falling slips, delivered using a split-belt treadmill. By measuring points of subjective equality over multiple perturbation sizes, we determined biases in people’s perception of risk associated with trips and slips. Both young and older adults perceived slips to be riskier than trips, but this perceptual bias did not differ between age groups. Understanding how people evaluate physical risk while walking is important for determining how a person’s prior experience influences their likelihood of making decisions that might put them at risk of falls.

While previous studies have not directly compared the perception of trips and slips, prior work has demonstrated that treadmill-generated backward-falling slips generate larger whole-body CoM displacements and quicker stepping responses compared to forward-falling trips [15]. People have also been reported to fall more frequently in response to slips versus trips of the same magnitude when performing perturbation-based training[36,37]. These results suggest that slips may generally be more difficult to recover from than trips. Consistent with these results, we found that the change in peak CoM velocity relative to unperturbed walking was higher during slips than trips. A strength of our analysis was that CoM velocity was only compared between trips and slips that were of equal magnitude in terms of acceleration and change in treadmill belt speed. Our results complement prior observations by establishing that people also perceive slips to be riskier.

Contrary to our hypothesis, the degree of sensitivity to perturbation direction was not different between our young and older adult participants, nor did the changes in CoM velocity in response to perturbations differ between groups. These results are contrary to previous work that has shown that older adults use more ineffective strategies and are less able to successfully recover from both trips and slips than young adults[38–40]. Although our older adults had poorer balance than our young adults, as measured by the MiniBESTest, both groups were similar with respect to self-reported physical activity, balance confidence, and walking speed. 84% of our older adults reported engaging in at least 150 minutes of exercise per week, which is the recommended amount of exercise according to the 2008 Physical Activity Guidelines for Americans[41]. This percentage of older adults engaged in exercise is much higher than that reported for the general population of older adults (27.3% - 44.3%)[42]. These findings suggest that our sample may not be representative of the general older adult population. While some older adults in our sample reported experiencing a fall in the previous year, this alone did not differentiate them from non-fallers in terms of sensitivity to perturbation direction, CoM velocity responses to perturbations and state anxiety. Future work should examine if older adults with lower balance confidence and physical activity or fitness levels exhibit greater sensitivity to perturbation direction, and if so, if this is then related to their self-reported fear of falling or fall history[43–45].

We found that the sensitivity to perturbation direction was stable across two days of testing such that those who found slips to be riskier on the first day also found them to be riskier on the second day. However, the degree of sensitivity was lower on the second day. Previous work has shown that both young and older adults are able to improve their reactive responses to trips and slips with perturbation training[46–48]. It is unclear whether this adaptation of reactive responses is associated with a reduction in perceived risk or balance threat posed by these losses of balance. Emotional arousal and anxiety have been shown to decrease with repeated exposure to the threat of standing perturbations[49]. Therefore, it is possible that our observation of reduced sensitivity to perturbation direction on the second day of testing may be due to a reduction in the perceived risk of slips due to repeated exposure on the first day. Future studies should assess emotional responses at various time points during protocols that require experiencing multiple gait perturbations to better understand the underlying mechanisms of any observed behavioral responses to the perturbations.

Several potential factors may lead to differences in people’s perceived risk associated with trips versus slips. First, if the two directions of perturbations elicit distinct emotional responses, these responses may inform an individual’s risk perception and choices. In behavioral economics, this is referred to as affect-based decision-making, where affect is the overall feeling of “good” or “bad” associated with a stimulus[12]. Anxiety in response to postural perturbations has been found to be correlated with self-reported perceptions of postural instability in both young[16,50] and older adults[50]. Therefore, we hypothesized that measures of state anxiety in response to slips and trips would correlate with people’s preferences between the two. However, we found no differences in self-reported state anxiety between the two perturbation types and between the two age groups. We also did not find a correlation between state anxiety and preference between trips and slips. This may be in part due to the limitations of self-reports themselves, as participants may use different frames of reference for their responses[51]. Therefore, some participants may have reported their anxiety in the context of falling such that if they did not fall, they reported low anxiety. Since our perturbations did not cause falls, this could have resulted in similar anxiety scores between trips and slips. This limitation can be addressed by approximating anxiety using an objective and instantaneous measure of arousal, such as electrodermal activity[52–54], which has previously been found to be sensitive to postural threat[17].

A second potential factor that may explain people’s preference between trips and slips is the actual physical disturbance of the body to the two perturbation directions. A study of perception thresholds of treadmill slips found that CoM velocity may be an important source of sensory information used by the nervous system to detect slips[35]. Therefore, we hypothesized that peak CoM velocity during trips and slips may be used as a source of sensory information that informs perceptions of severity and, thus, may explain people’s preferences. However, we found that relative changes in peak CoM velocity from baseline during trips and slips did not correlate with our measure of preference. It is possible that a combination of proprioceptive signals related to foot, lower limb, and head kinematics may better explain people’s preferences. A final consideration is that in our experimental setup, handlebars were available towards the front of the treadmill. While participants were verbally discouraged from constant use of this external support, some did still use them in anticipation of perturbations. Therefore, the measure CoM velocity following perturbations was occasionally confounded by the use of support bars. Future studies can either remove support bars, if possible, or record their use so that they can be accounted for.

## Conclusion

Both young and older adults perceive slips to be riskier than trips, although the underlying reasons for this preference remain to be understood. Measuring the degree of preference for different perturbation types, along with a person’s ability to recover from them, may help identify activities that are most likely to cause falls. This information could be used to personalize training programs in accordance with an individual’s risk perception and balance capacity. Subsequent studies should also examine how the characteristics of experienced losses of balance inform subsequent behavioral decisions. For example, epidemiological data on the prevalence of falls is often collapsed across the causes or perceived causes of falls such that slips and trips are grouped together[7,9]. In future studies, it may be valuable to consider the type of loss of balance when characterizing falls and then determine if this information predicts subsequent changes in behavior.

